# Comparative analyses of gene expression in common marmoset and human pluripotent stem cells (PSCs) identify factors enhancing homologous recombination efficiency in the *HPRT* locus of human PSCs

**DOI:** 10.1101/2021.04.05.438539

**Authors:** Sho Yoshimatsu, Mayutaka Nakajima, Tsukasa Sanosaka, Tsukika Sato, Hideyuki Okano

## Abstract

A previous study assessing the efficiency of the genome editing technology CRISPR-Cas9 for knock-in gene targeting in common marmoset (marmoset; *Callithrix jacchus*) embryonic stem cells (ESCs) unexpectedly identified innately enhanced homologous recombination (HR) activity in marmoset ESCs (cmESCs). Here, we compared gene expression in marmoset and human pluripotent stem cells (PSCs) using transcriptomic and quantitative PCR (qPCR) analyses and found that five HR-related genes (*BRCA1, BRCA2, RAD51C, RAD51D* and *RAD51*) were upregulated in marmoset cells. Four of these upregulated genes enhanced HR efficiency with CRISPR-Cas9 in human pluripotent stem cells. Thus, the present study provides a novel insight into species-specific mechanisms for the choice of DNA repair pathways.

## 1. Introduction

Repairing DNA double strand breaks (DSBs) is indispensable for maintaining genomic stability (Schipler and Iliakis, 2013). From prokaryotes to eukaryotes, conserved repair pathways for DSBs have been identified: nonhomologous end-joining (NHEJ); homology-directed repair (HDR), including homologous recombination (HR); and microhomology-mediated end-joining (MMEJ), also known as alternative nonhomologous end-joining. The choice of which repair pathways are employed relies on a variety of factors, including DSB complexity, cell type, cell cycle, and species (Buerstedde and Takeda, 1991; Schipler and Iliakis, 2013; Scully et al., 2019; Sung and Klein, 2006).

The genome editing CRISPR-Cas9 technology (Jinek et al., 2012) relies on endogenous pathways for repairing DSBs in cells (Cong et al., 2013). Although the combination of CRISPR-Cas9 and double-stranded DNA (dsDNA) donor enables precisely targeted integration or deletion of long sequences by HDR, the efficiency of this process is limited by competing DSB repair pathways, mainly NHEJ. Previous studies have demonstrated that the efficiency of Cas9-meditated HDR can be modified through use of small molecules that stimulate the HDR-related factor RAD51 (Pinder et al., 2015; Song et al., 2016) or that inhibit the NHEJ-related factor LIG4 (Chu et al., 2015; Maruyama et al., 2015). Additionally, modification of Cas9 by fusion with DSB repair-related factors such as RAD52, CtIP, MRE11A (Charpentier et al., 2018; Shao et al., 2017; Tran et al., 2019), or a dominant negative mutant of P53BP1 (Jayavaradhan et al., 2019) can improve HDR efficiency. Also, overexpression of several DSB-related genes, including *RAD52* (Shao et al., 2017) and *RAD18* (Nambiar et al., 2019), reportedly contributed to enhancement of Cas9-mediated HDR efficiency.

Human pluripotent stem cells (hPSCs), including human embryonic stem cells (hESCs) and induced pluripotent stem cells (hiPSCs), hold considerable promise for a wide range of applications in the fields of regenerative medicine and stem cell biology (Okano and Sipp, 2020; Okano and Yamanaka, 2014; Yamanaka, 2012; Yamanaka, 2020). Moreover, knock-in gene targeting (KI) mediated by HR in PSCs offers a powerful approach to the directed differentiation of PSCs (Irie et al., 2015; Sasaki et al., 2015; Sybirna et al., 2020; Yoshimatsu et al., 2019b; Zwaka and Thomson, 2003), disease modeling (Fujimori et al., 2018; Ichiyanagi et al., 2016; Nakamura et al., 2019), and gene therapy (Nakamoto et al., 2018; Ou et al., 2016; Simara et al., 2013). However, although the application of CRISPR-Cas9 greatly facilitates HR-mediated KI in hPSCs, there is still room for further improvement, such as enhancing the ratio of homologous recombinants to total drug-resistant clones including non-recombinants (genomic integration of a drug resistance cassette by NHEJ-mediated random integration; hereinafter HR ratio). In hPSCs, several studies have shown that even with use of site-specific nucleases, such as zinc finger nucleases (ZFNs), transcription activator-like effector nucleases (TALENs), and CRISPR-Cas9, the HR ratio is less than 50% and generally closer to 30% (Hockemeyer et al., 2009; Hockemeyer et al., 2011; Sebastiano et al., 2011; Takayama et al., 2017).

In a previous study (Yoshimatsu et al., 2019a), we showed that common marmoset (marmoset; *Callithrix jacchus*) ESCs (cmESCs) harbor an unusual facility for DSB repair, which is characterized by a high HR ratio. In KI experiments using a *PLP1-EGFP* vector targeting the 1st exon of the *PLP1* gene, we found rates of homologous recombinants of 90% with the use of CRISPR-Cas9 and 80% without its use (Yoshimatsu et al., 2019a). Moreover, we observed high HR ratios in a variety of other loci, including *ACTB, PLP1* (targeting the 2nd, 5th, and 6th exons), *FOXP2, PRDM1, DPPA3* (Yoshimatsu et al., 2019a; Yoshimatsu et al., 2020; Yoshimatsu et al., 2019b), and *NANOS3* (Figure S1). On the basis of these results, we have now explored possible mechanisms for the HR-biased DSB repair in cmESCs. Through use of interspecies analyses of gene expression, we have identified factors that enhance the HR ratio with CRISPR-Cas9 in hiPSCs by ectopic overexpression.

## 2. Materials and Methods

### 2.1 Ethical statement

Recombinant DNA experiments were approved by the Recombinant DNA Experiment Safety Committee of Keio University (approval number: 27-023 and 27-034).

### 2.2 Vector construction

The human *HPRT* targeting vector HPRT-TV was constructed using the GKI-2.0 system (Yoshimatsu et al., 2019b). In brief, based on sequence information of the human X chromosome from the NCBI genome browser (GRCh38.p13, Chromosome X - NC_000023.11: 134460165..134500668), the 5’ homology arm (3.0 kb) and 3’ homology arm (12.8 kb), which are around the 2nd intron of the gene, were subcloned respectively into pDONR-P3P1r (Addgene #141014) and pDONR-P2rP4 (Addgene #141015) using the Gateway cloning system (BP reaction). The targeting vector containing a *PGK-Neo* drug-resistance cassette flanked by the *HPRT* homology arms was constructed by Gateway LR reaction using pENTR-L1-PGK-Neo-L2 (Addgene #141007), pUC-DEST-R3R4(R) (Addgene #141010) and the Gateway LR Clonase II Enzyme Mix (Thermo Fisher Scientific).

A marmoset *NANOS3-Venus* targeting vector was constructed using the GKI-3.0 system (Yoshimatsu et al., 2019b), which contained the monomeric *Venus* (an improved version of *yellow fluorescent protein*) (Kremers et al., 2006; Nagai et al., 2002) encoding gene and a puromycin/thymidine kinase positive/negative selection cassette flanked by homology arms of the marmoset *NANOS3* gene. In brief, based on information from the NCBI genome browser (GCF_000004665.1, Chromosome 22 - NC_013917.1: 13114410..13117680), the 5’ homology arm (540 bp) and 3’ homology arm (542 bp) were cloned respectively into pENTR2-L3-SfoI-Venus-PBL-R1 (Addgene #141019) and pDONR-P2rP4 (Addgene #141015) using the Seamless Cloning method (Thermo Fisher Scientific). The targeting vector containing a *PGK-PuroTK* cassette flanked by the corresponding homology arms was constructed by Gateway LR reaction using pENTR-L1-PGK-PuroTK-L2 (Addgene #141005), pUC-DEST-R3R4(R) (Addgene #141010) the Gateway LR Clonase II Enzyme Mix.

PX459 was used as a Cas9/gRNA vector (Ran et al., 2013); the vector contained either the human *HPRT* sgRNA sequence TGTTTCAATGAGAGCATTAC or the marmoset *NANOS3* sgRNA sequence TCCTTCCATGTCCACCTAGG.

*RAD51, RAD51C*, and *RAD51D* overexpression vectors were constructed using a pDONR-R4-CAG-pA-L2 backbone (kindly provided by Takefumi Sone at Keio University). *BRCA1* and *BRCA2* overexpression vectors were constructed using pcBRCA1-385 (a gift from Lawrence Brody, Addgene plasmid #61586) and pcDNA3 236HSC WT (BRCA2) (a gift from Mien-Chie Hung, Addgene plasmid #16246) (Wang et al., 2002), respectively, as backbones. SV40 promoter-driven neomycin-resistance cassettes in pcBRCA1-385 and pcDNA3 236HSC WT (BRCA2) were truncated and destroyed by *Sma*I and *Sfo*I digestion and subsequent ligation.

### 2.3 Cell culture, transfection and drug selection

Three cmESC lines were used: No. 40 (CMES40) and No. 20 (CMES20) (Sasaki et al., 2005), and DSY127 (Yoshimatsu et al., 2020); all three cell lines were cultured as described previously (Yoshimatsu et al., 2019a). Vector transfection into cmESCs and puromycin selection were performed as described previously (Yoshimatsu et al., 2019a).

Three human iPSC lines, 201B7 (Takahashi et al., 2007), WD39 (Imaizumi et al., 2012) and etKA4 (Matsumoto et al., 2016), and a human ESC line KhES-1 (Okita et al., 2011) were used; the cell lines were cultured as described previously (Matsumoto et al., 2016). In brief, PSCs were cultured on mitomycin-C-treated G418-resistant SNL76/7 feeder cells (McMahon and Bradley, 1990) in ESM under 20% O_2_ and 5% CO_2_ at 37°C. ESM consisted of 1× DMEM/F12 (Thermo Fisher Scientific) supplemented with 20% Knockout Serum Replacement (Thermo Fisher Scientific), 0.1 mM MEM Non-Essential Amino Acids Solution (Nacalai Tesque), 1 mM L-glutamine (L-glu; Nacalai Tesque), 0.1 mM 2-mercaptoethanol (2-ME; Sigma), 100 U/ml penicillin and 100 µg/ml streptomycin sulfate (Nacalai Tesque) and 4 ng/ml basic fibroblast growth factor (Peprotech).

For passaging, cells were pre-treated with 10 μM Rho-associated coiled-coil-containing protein kinase inhibitor Y-27632 (Wako) in ESM at 37°C for an hour. The cells were then incubated in CTK solution (Reprocell) at 37°C for 30 sec, mechanically separated from feeder cells, and dissociated by gentle pipetting. The isolated cells were plated onto new feeder cells in ESM supplemented with 10 μM Y-27632. Twenty-four hours later, Y-27632 was removed from the medium. Medium change was performed daily. Prior to the seeding of hESCs/iPSCs, feeder cells were seeded onto a gelatin-coated 100 mm cell culture dish in Dulbecco’s modified Eagle’s medium (Thermo Fisher Scientific) supplemented with 10% inactivated fetal bovine serum (Thermo Fisher Scientific).

For transfection (Day 0), we used the NEPA21 Super Electroporator (Nepagene) as described previously (Yoshimatsu et al., 2019b). Ten μg of HPRT-TV and 5 μg each of the Cas9/gRNA and overexpression vectors were diluted in 100 μl of OPTI-MEM (Thermo Fisher Scientific). The etKA4 hiPSCs (>1 × 10^7^ cells) were suspended in the solution and subjected to electroporation. Transfected cells were plated onto new mitomycin-C-treated G418-resistant SNL76/7 feeder cells on a 100-mm cell culture dish in ESM supplemented with 10 μM Y-27632. On Day 2, selection was initiated by adding 100 ng/ml G418 (an analog of neomycin; Sigma) to the ESM; selection was performed for six days. On Day 8, the concentration of G418 was doubled, and selection was continued for an additional three days. On Day 11, the number of G418-resistant colonies was counted. The cells were then subjected to further selection in 10 μM 6-thioguanine (6TG) for five days. On Day 16, the number of 6TG-resistant colonies was counted. Experimental data was only included when at least 80 G418-resistant colonies survived.

### 2.4 Genotyping

Southern blotting was performed as described previously (Yoshimatsu et al., 2019a) using the digoxigenin (DIG) probe system. Genomic DNA samples were digested with *Bgl*II and *Eco*RV by overnight 37°C incubation. We used the PCR DIG Probe Synthesis Kit (Roche) for DIG-labeled probe production. The human *HPRT*-specific probe (492 bp) was amplified from human genomic DNA using the primers TGCATATCTGGGATGAACTCTGG and AAATGGGACATTTGTGTGTCACC. Molecular sizes were confirmed using DIG-labeled DNA Molecular Weight Marker II (λDNA with *Hin*d III digestion) (Sigma; #11218590910). Genotyping PCR and DNA sequencing analysis (Figure S1) were performed as described previously (Yoshimatsu et al., 2019b).

### 2.5 Quantitative reverse-transcription PCR (qPCR)

RNA extraction, reverse transcription, and PCR were performed as described previously (Yoshimatsu et al., 2019a). Three biological and technical repetitions of the qPCR analyses were performed. Quantification was performed using the relative standard curve method and endogenous expression of *glyceraldehyde 3-phosphate dehydrogenase* (*GAPDH*) was used as an internal control. Primers were newly or previously designed to amplify both human and marmoset cDNA sequences; however, they did not amplify murine cDNA sequences to avoid contamination due to the use of MEFs for PSC culture (Yaglom et al., 2014; Yoshimatsu et al., 2019a). The following primers were used: *GAPDH*-forward (GCACCGTCAAGGCTGAGAAC), *GAPDH*-reverse (TGGTGAAGACGCCAGTGGA), *RAD51*-forward (GTCACCTGCCAGCTTCCCATT), *RAD51*-reverse (AGCAGCCGTTCTGGCCTAAAG), *RAD51C*-forward (CGCTGTCGTGACTACACAGAGT), *RAD51C*-reverse (AGGCTGATCATTTGCTGGGCT), *RAD51D*-forward (GGTGCTGCTGGCTCAGTTCT), *RAD51D*-reverse (CGCTACCTGGGCCTCCTACA), *BRCA1*-forward (ACCCGAGAGTGGGTGTTGGA), *BRCA1*-reverse (GCTGTGGGGGATCTGGGGTA), *BRCA2*-forward (TGGGCTCTCCTGATGCCTGTA) and *BRCA2*-reverse (GTATACCAGCGAGCAGGCCG).

### 2.6 RNA-seq analysis

Transcriptome data was obtained from cmESCs as described previously (GSE138944) (Shiozawa et al., 2020). We used the deposited RNA-seq data from human ESCs/iPSCs (GSE53096) (Ma et al., 2014) as a reference. Marmoset mRNA was sequenced on an Illumina HiSeq2500 and the obtained nucleotide sequences were mapped against the *Callithrix jacchus* genome (Callithrix_jacchus_cj1700_1.1; https://www.ncbi.nlm.nih.gov/assembly/GCF_009663435.1/) by *STAR* (ver.2.5.3a). The number of mapped reads was counted by *featureCounts* (1.5.2) and simultaneously normalized by the TMM method in the *edgeR* package in *R* (Robinson et al., 2010). The normalized expression levels processed to log2 and z-scoring were visualized using the pheatmap library. In the statistical analysis, Welch’s *t-*test was performed between normalized gene expression levels of human samples and marmoset samples, and the resulting *p* values were processed with Bonferroni correction to obtain the adjusted *p* value. In the present study, adjusted *p* values less than 0.05 were defined as significant; adjusted *p* values processed to -log10 were visualized on the vertical axis by the *ggplot2* library in *R*.

### 2.7 Western blotting

Western blotting was performed using the Wes - Automated Western Blots with Simple Western (ProteinSimple) according to the manufacturer’s introductions. As primary antibodies, we used polyclonal Rad51 H-92 antibody (1:50 dilution; sc-8349; Santa Cruz) and monoclonal α-tubulin antibody (1:25000 dilution; T9026; Sigma), which was used as an internal control for RAD51 protein quantification. ImageJ software was used to quantify the intensities of Rad51 (37 kDa) and α-tubulin (50 kDa) bands, and then RAD51 expression was normalized against α-tubulin expression.

### 2.8 Statistical analysis

All data in this study are expressed as means ± S.D. Statistically significant differences were determined using Welch’s *t*-test; *p* values < 0.05 are designated by *; *p* values < 0.01 are designated by **, and are interpreted as statistically significant.

## 3. Results

### 3.1 Comparative transcriptomic analysis of human and marmoset PSCs

Previously, we reported that cmESCs have innately high HR activity (Yoshimatsu et al., 2019a). In particular, we observed extraordinarily high HR ratios for the 1st exon of *PLP1* in a targeting experiment (92.3% with CRISPR-Cas9 against 88.6% without its use). We also observed high HR ratios using CRISPR-Cas9 in other loci, such as *ACTB, PLP1* (targeting the 2nd, 5th, and 6th exons), *FOXP2, PRDM1, DPPA3*, and *NANOS3* (Yoshimatsu et al., 2019a; Yoshimatsu et al., 2020; Yoshimatsu et al., 2019b) see Figure S1). Here, we investigated possible factors that might underlie this phenomenon.

Initially, we investigated HR- and NHEJ-related gene expression in human and marmoset PSCs. In hPSCs, several studies have shown that the HR ratio is less than 50%, generally around 30%, even with use of site-specific nucleases such as ZFN, TALEN, and CRISPR-Cas9 (Hockemeyer et al., 2009; Hockemeyer et al., 2011; Sebastiano et al., 2011; Takayama et al., 2017). We used RNA-seq data of hESCs/iPSCs deposited in databases (Ma et al., 2014) to compare gene expression with cmESCs as described previously (Shiozawa et al., 2020). We merged and normalized PSC RNA-seq data derived from both species.

HR- and NHEJ-related genes that can be used for comparing fold changes between hPSCs and cmESCs have been listed by hsa03440 and ko03450 in KEGG (https://www.genome.jp/kegg/pathway.html). Some related genes are absent from the gene lists due to the incomplete gene assembly of the latest version of the marmoset genome (Callithrix_jacchus_cj1700_1.1). Here, we analyzed 37 HR-related and 12 NHEJ-related genes (Figures 1A−D and S2−3). As summarized in Supplementary Data 1, eleven genes (*RAD51D, BRCA1, BRCA2, BABAM1, RAD51, RAD51C, POLD2, RAD51B, MUS81, POLD1* and *XRCC3*) were significantly up-regulated in cmESCs (Figure 1A), whereas eleven others (*RPA2, ATM, XRCC2, SSBP1, RPA3, PALB2, BRIP1, NBN, RAD54B, UMC1* and *BRCC3*) were down-regulated (Figure 1B). With regard to NHEJ-related genes, six genes (*XRCC6, PRKDC, POLM, XRCC5, RAD50* and *DNTT*) were significantly up-regulated in cmESCs (Figure 1C), and five others (*FEN1, DOLRE1C, NHEJ1, XRCC4* and *POLL*) were down-regulated (Figure 1D). In light of the high HR activity in cmESCs, we then focused on HR-related genes that showed increased expression in cmESCs compared to hPSCs. To validate the results of the transcriptome analysis, we performed an interspecies qPCR analysis using primer sets specifically designed for these human and marmoset genes. We also designed primers specific for human and marmoset *GAPDH* for normalization. Preliminary screens using the designed primers showed that an interspecies comparison of expressions was not feasible for several genes (*BABAM1* and *POLD1/2*) owing to a lack of accuracy based on the post-qPCR melt curve analysis (data not shown).

**Figure 1.**
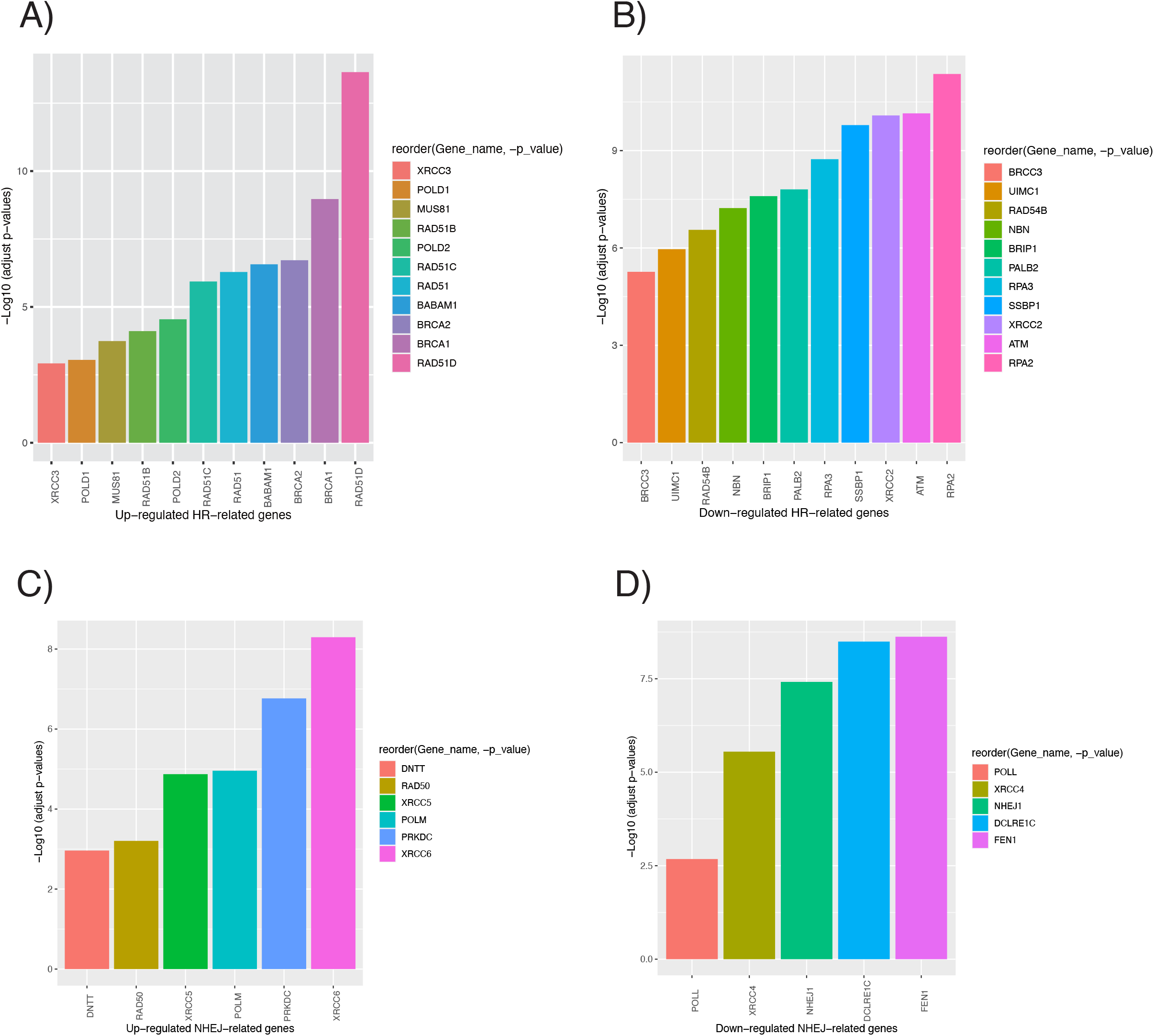
Differential expression of DSB repair genes in human and marmoset PSCs. (**A−B**) HR-related genes that are significantly upregulated or downregulated in cmESCs compared to those in hESCs/iPSCs. Gene groups are referenced to KEGG hsa03440. Y-axis shows adjusted p-values after -log10 treatment. (**C−D**) NHEJ-related genes showing significantly higher or lower expression levels in cmESCs compared to hESCs/iPSCs. Gene groups refer to KEGG ko03450. Y-axis shows the adjusted p-value after -log10 treatment.

By qPCR using total RNAs from three cmESC lines (No. 40, No. 20, and DSY127) and four hPSC lines (201B7, WD39, KhES-1, and etKA4), we confirmed the significantly higher expression of *RAD51D, BRCA1, BRCA2, RAD51C*, and *RAD51* in cmESCs compared to hPSCs (Figure 2). In particular, *RAD51C* and *RAD51D* expression in cmESCs was approximately 10 and 7 times higher, respectively, than those of hPSCs (Figure 2, top). Quantitative western blotting confirmed the high RAD51 expression in cmESCs at the protein level (Figures S4A−B).

**Figure 2.**
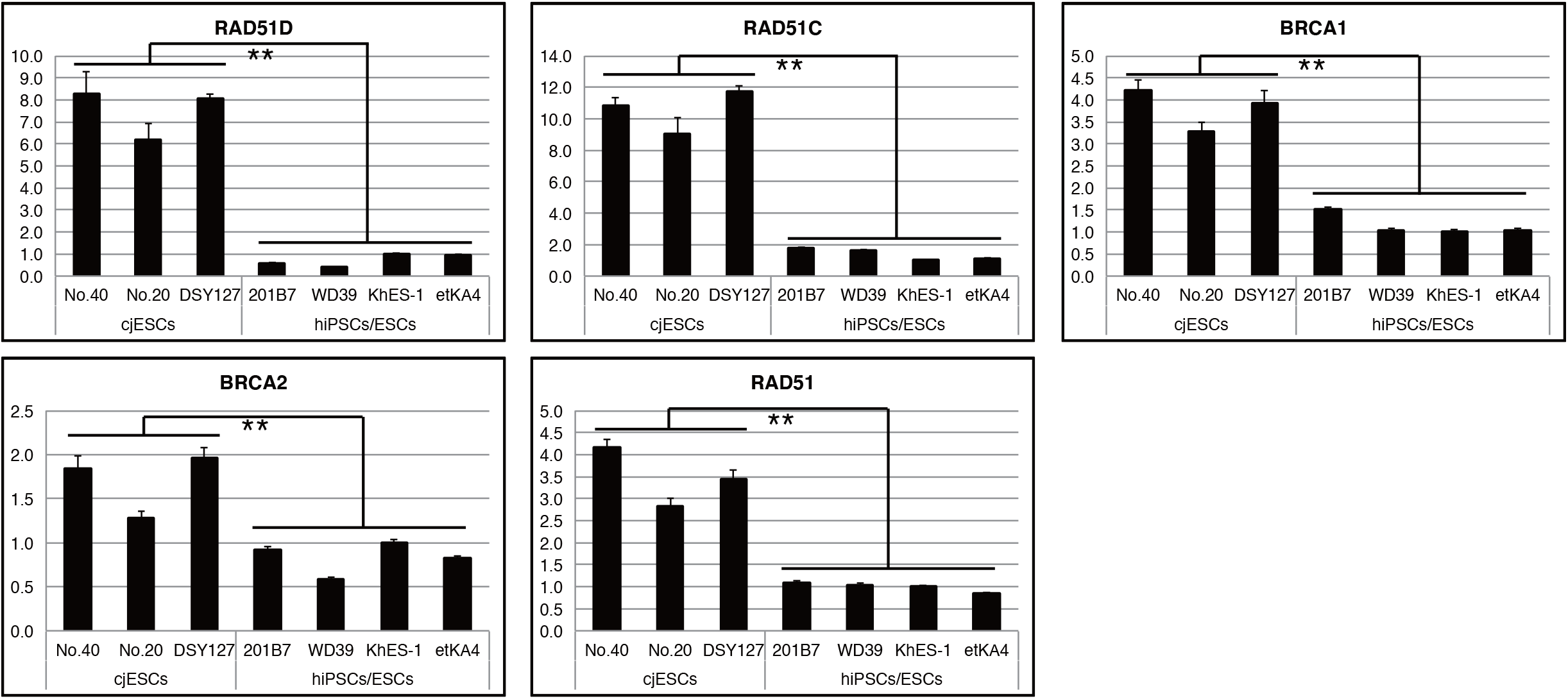
Interspecies qPCR analysis for *RAD51C, RAD51D, BRCA1, BRCA2* and *RAD51*. RQ values of each human/marmoset sample (biological and technical triplicates) were used for the statistical significance tests.

### 3.2 Enhancement of the HR/RI ratio with CRISPR-Cas9 in hiPSCs by overexpression of the defined factors

To explore the effect of high expression of the five HR-related genes (*RAD51D, RAD51C, BRCA2, RAD51*, and *BRCA1*) in cmESCs, we induced overexpression of the genes in a gene targeting experiment with *hypoxanthine-guanine phosphoribosyltransferase* (*HPRT*) in the male hiPSC line etKA4 (Matsumoto et al., 2016). The HPRT protein catalyzes the salvage pathway, synthesizing inosine monophosphate and guanosine monophosphate from hypoxanthine and guanine, respectively (Torres and Puig, 2007). HPRT deficiency results in the loss of susceptibility for 6TG, a toxic analog of guanine (Sharp et al., 1973; Wahl et al., 1975). We selected the HPRT targeting system as it has been frequently used to assess the HR ratio in mammalian male PSCs (Meek et al., 2010; Thomas and Capecchi, 1987; Zwaka and Thomson, 2003).

We constructed a knock-in/knock-out system for the human *HPRT* gene (Materials and Methods). Following KI, the 2nd exon of *HPRT* was completely replaced with a PGK-Neo cassette, which resulted in the loss of functional mRNA expression from the *HPRT*^Neo^ allele (Figure 3A). Initial G418 selection (both homologous recombinants and non-recombinants survived) and subsequent 6TG selection (only homologous recombinants survived) enabled robust quantification of the HR ratio without the necessity of genotyping individual clones (Figure 3B). In addition, we constructed a PX459-based Cas9/gRNA vector (Ran et al., 2013) containing the sgRNA sequence for the 2nd intron of *HPRT*, which did not recognize HPRT-TV.

**Figure 3.**
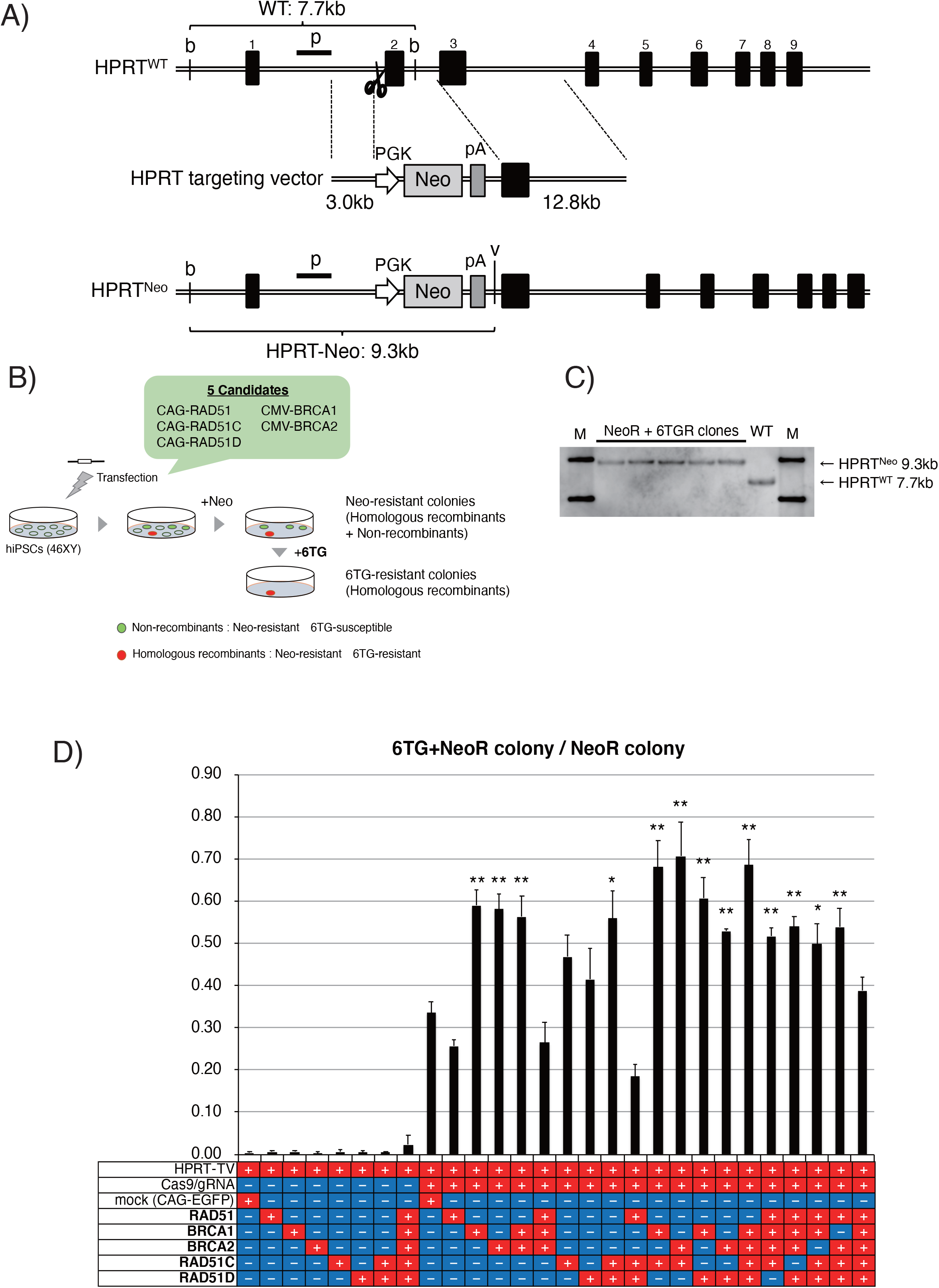
*HPRT* targeting in hiPSCs. (**A**) Graphical schematics of the wild-type human *HPRT* locus (*HPRT*^WT^; top), *HPRT* targeting vector (middle), and recombinant *HPRT* locus (*HPRT*^Neo^; bottom). Black boxes indicate endogenous exons of the *HPRT* gene (upper numbers indicate each exon number). *PGK*, mouse *phosphoglycerate kinase 1* promoter; *Neo, neomycin resistance gene*; *pA, polyadenylation signal sequence*; b, *Bgl*II recognition site; v, *Eco*RV recognition site; p, detection site of the DIG-labelled probe for Southern blotting analysis. (**B**) Graphical schematic of the HPRT targeting experiment using hiPSCs. *CAG, CAG promoter*; *CMV, human cytomegalovirus immediate early enhancer and promoter*. (**C**) Southern blotting analysis of genomic DNA derived from G418/6TG-double-resistant five clones and wild-type (WT) hiPSCs. M, DNA marker (λDNA with *Hin*d III digestion). (**D**) Resultant HR/RI ratios in the HPRT targeting experiments. Values were calculated as 6TG+NeoR colony number / NeoR colony number. Asterisks indicate statistical significance in comparisons of Cas9/gRNA (+) samples of control versus Cas9/gRNA (+) mock (+).

As it is possible that the genomic cleavage of the *HPRT* 2nd intron after transfection of the Cas9/gRNA vector and subsequent NHEJ or MMEJ-mediated introduction of small/large deletion could produce an undesired knock-out allele, we initially tested transfection of only the Cas9/gRNA vector into the etKA4 hiPSCs. In this initial test, no 6TG-resistant colonies were obtained from 1 × 10^7^ transfected cells (n = 3), showing that the NHEJ or MMEJ-mediated deletion in the intronic region has a negligible effect with regard to the assessment of the HR ratio in the hiPSCs.

After co-transfection of the Cas9/gRNA vector and HPRT-TV, and serial G418 and 6TG selection, we confirmed that all analyzed G418 and 6TG-resistant (NeoR+6TGR) clones were hemizygous *HPRT*^Neo^ recombinants by Southern blotting (Figure 3C).

Since we wished to evaluate the effects of single/multiple overexpression of the five genes on HR ratios (Figure 3B), we used a CAG-EGFP vector (Shiozawa et al., 2020) as a mock control for the overexpression vectors. Initially, we transfected HPRT-TV and each overexpression vector (or mock) and quantified HR ratios without CRISPR-Cas9. None of the attempts at single/multiple overexpression of the five genes enhanced HR ratios significantly (Figure 3D; mock, 0.0033 ± 0.0018; +RAD51, 0.0060 ± 0.0040; +BRCA1, 0.0060 ± 0.0030; +BRCA2, 0.0026 ± 0.0043; +RAD51C, 0.0052 ± 0.0052; +RAD51D, 0.0048 ± 0.0035; +RAD51C/D, 0.0045 ± 0.0029; +BRCA1/2&RAD51C/D, 0.023 ± 0.023; n ≧ 4).

Next, we quantified HR ratios with use of CRISPR-Cas9. When the mock vector was transfected with HRPT-TV and the Cas9/gRNA vector, an HR ratio of approximately 30% was achieved (Figure 3D; 0.336 ± 0.026, n = 4); this HR ratio is comparable with previous results using hPSCs (Takayama et al., 2017). Overexpression of only *RAD51* did not result in a significant enhancement of the HR ratio (0.256 ± 0.163, n = 3).

In comparison to use of the mock, single overexpression of *BRCA1* or *BRCA2* surprisingly resulted in a significant enhancement of the HR ratio (Figure 3D; +BRCA1, 0.590 ± 0.036; +BRCA2, 0.582 ± 0.034; n = 3). In addition, co-overexpression of *BRCA1* and *BRCA2* also enhanced the HR ratio (0.563 ± 0.049, n = 3), while co-overexpression of *BRCA1/2* and *RAD51* did not produce an enhanced HR ratio (0.265 ± 0.049, n = 3).

Single overexpression of *RAD51C* or *RAD51D* did not result in a significant enhancement of the HR ratio (Figure 3D; +RAD51C, 0.468 ± 0.051; +RAD51D, 0.414 ± 0.073; n = 4). However, when the two genes were co-overexpressed, we observed an enhanced HR ratio (0.560 ± 0.065, n = 3); however, this effect was not observed when *RAD51* was also co-overexpressed (0.185 ± 0.03, n = 3).

We tested multiple sets of overexpression of the five genes (Figure 3D). Except when RAD51 was co-transfected with the other four genes (0.388 ± 0.032, n = 3), co-overexpression of *RAD51C/D* and *BRCA1/2* resulted in significant enhancement of HR ratios: +RAD51C and BRCA1, 0.681 ± 0.062; +RAD51C and BRCA2, 0.706 ± 0.080; +RAD51D and BRCA1, 0.607 ± 0.048; +RAD51D and BRCA2, 0.529 ± 0.006; +RAD51C and BRCA1/2, 0.516 ± 0.021; +RAD51D and BRCA1/2, 0.540 ± 0.024; +RAD51C/D and BRCA1, 0.498 ± 0.048; +RAD51C/D and BRCA2, 0.539 ± 0.044 (n = 3 for all combinations). Finally, we demonstrated that co-overexpression of the factors *BRCA1, BRCA2, RAD51C*, and *RAD51D* resulted in a significant enhancement of the HR ratio (0.686 ± 0.061, n = 3).

## 4. Discussion

In this study, through comparative analyses of gene expression in human and marmoset PSCs, we have identified four genes (*RAD51C, RAD51D, BRCA1*, and *BRCA2*) whose single/multiple overexpression increased HR ratios in hiPSCs. Intriguingly, we also observed that overexpression of *RAD51* did not enhance the HR ratio in hiPSCs; an alternative explanation is that *RAD51* overexpression cancelled the effect of the enhancement of the HR ratio by the other four genes. In Supplementary Discussion, we further discuss how these genes involve in the HR machinery, and the discrepancy of the effect of *RAD51* overexpression from part of previous studies. Results presented here also suggests the possibility of a vice versa effect. In fact, we demonstrated the overexpression of four factors (*RAD51C, RAD51D, BRCA1*, and *BRCA2*), which were highly expressed in cmESCs, contributed to the enhancement of HR ratios in hPSCs. Thus, it is also possible that overexpression of several NHEJ factors, which were lowly expressed in cmESCs (including *FEN1, DCLRE1C* (*Artemis*), *NHEJ1*, and *XRCC4*), may contribute to decreased HR ratios in hPSCs. Further analyses are required to evaluate the robust effects of the four HR factors, such as effects on KI in other loci, and in different cell lines and species; nevertheless, our investigation has demonstrated that overexpression of these factors may ameliorate the HR ratio with CRISPR-Cas9 in hiPSCs.

In clinical settings, although knock-in technology in donor PSCs is beneficial due to the highly customizability, constitutive overexpression of exogenous gene(s) may impose potential risks including tumorigenicity and genome instability. In this context, critical time window of HR-factor overexpression for increasing HR ratios should be assessed in further studies.

## Author Contributions

Conceptualization, S.Y. and H.O.; methodology, S.Y.; software, S.Y., M.N. and T.Sanosaka ; validation, S.Y., M.N. and T.Sato; formal analysis, S.Y. and M.N.; investigation, S.Y., M.N. and T.Sanosaka; data curation, S.Y. and M.N.; writing—original draft preparation, S.Y.; writing—review and editing, S.Y., M.N. and H.O.; visualization, S.Y.; supervision, H.O.; project administration, H.O.; funding acquisition, H.O. and S.Y. (see the Funding section).

## Acknowledgments

We greatly thank Drs. Takashi Sasaki (Keio University) for kind support and technical advice on RNA-seq analysis, Ms. Kanae Ohtsu (Keio University) for technical support, Dr. Kent Imaizumi (Keio University) and Mr. Keisuke Oda (Ono Pharmaceutical Co., Ltd.) for kindly providing hPSC samples, Drs. Mitsuru Ishikawa (Keio University) and Takefumi Sone (Keio Univeristy) for technical advice on hPSC experiments, and Dr. Seiji Shiozawa (Keio University) for valuable comments. We also thank all the laboratory members of H.O. for their encouragement and generous support for this study.

---

**Figure S1.**
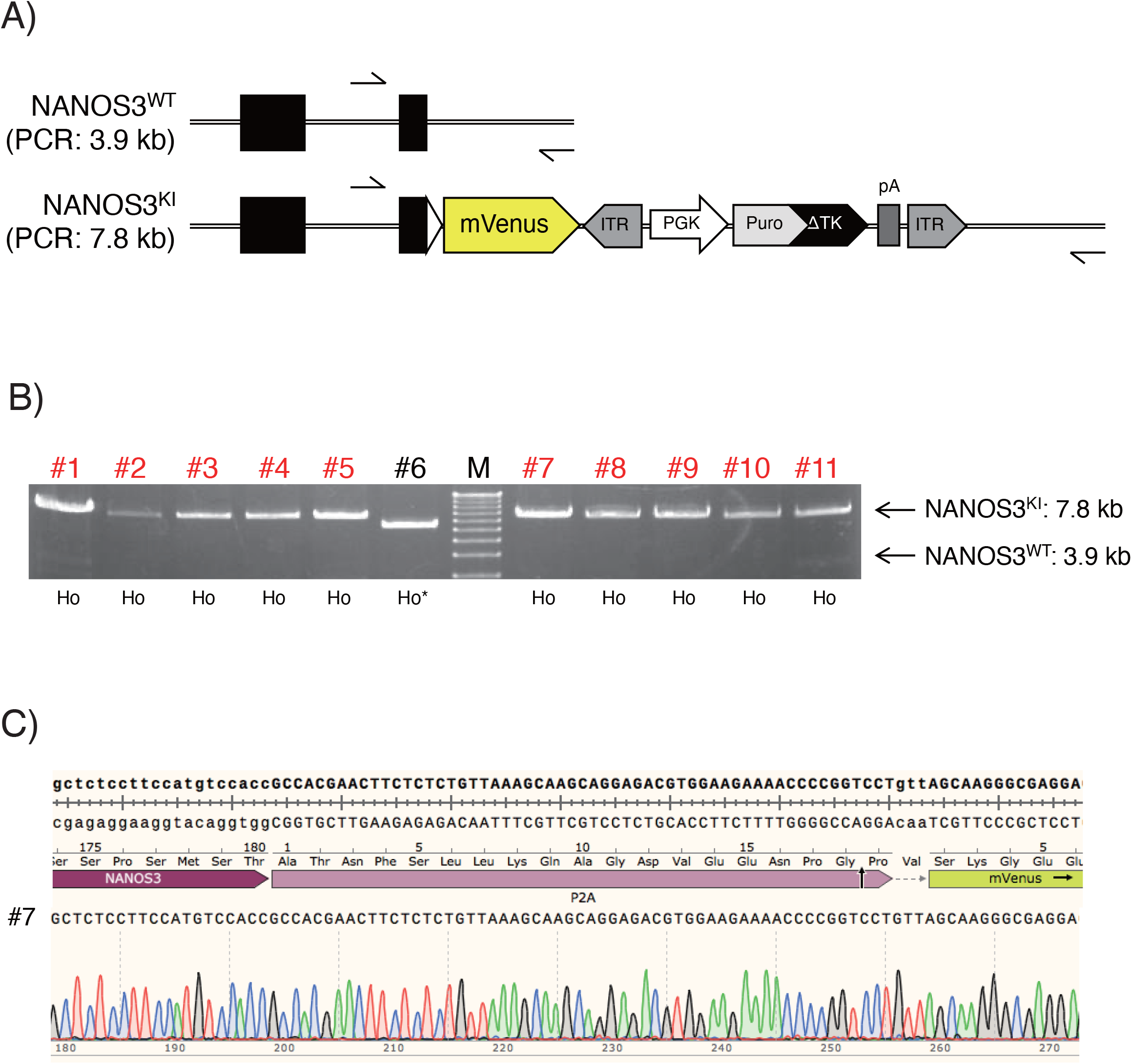
Gene targeting in the marmoset *NANOS3* gene locus. (**A**) Graphical schematics of the wild-type marmoset *NANOS3* locus (NANOS3^WT^; top), recombinant *NANOS3* locus (NANOS3^KI^; bottom). Black boxes and arrows indicate endogenous exons of the *NANOS3* gene and primer binding sites for genotyping PCR. White triangles indicate self-cleaving porcine teschovirus-1 2A sequence (P2A). ITR, excisable *Piggybac transposase* inverted terminal sequence; *Puro ΔTK, puromycin resistance gene* fused to the N-terminus truncated *thymidine kinase-1*. (**B**) Genotyping PCR analysis of puromycin-resistant cmESC clones following transfection of the *NANOS3-Venus* targeting vector and Cas9/gRNA vector. Ho, homozygous KI. All analyzed (eleven) clones harbored homozygous *NANOS3*^KI^ alleles, although clone #6 (Ho*) was homozygous for aberrant (truncated) recombinant alleles. A 1 Kb Plus DNA Ladder (Thermo Fisher Scientific) was used as a DNA marker. (**C**) DNA sequencing analysis of the recombinant alleles of clone #7. *P2A-Venus* was precisely integrated into the direct downstream region of the *NANOS3* coding sequence.

**Figure S2.**
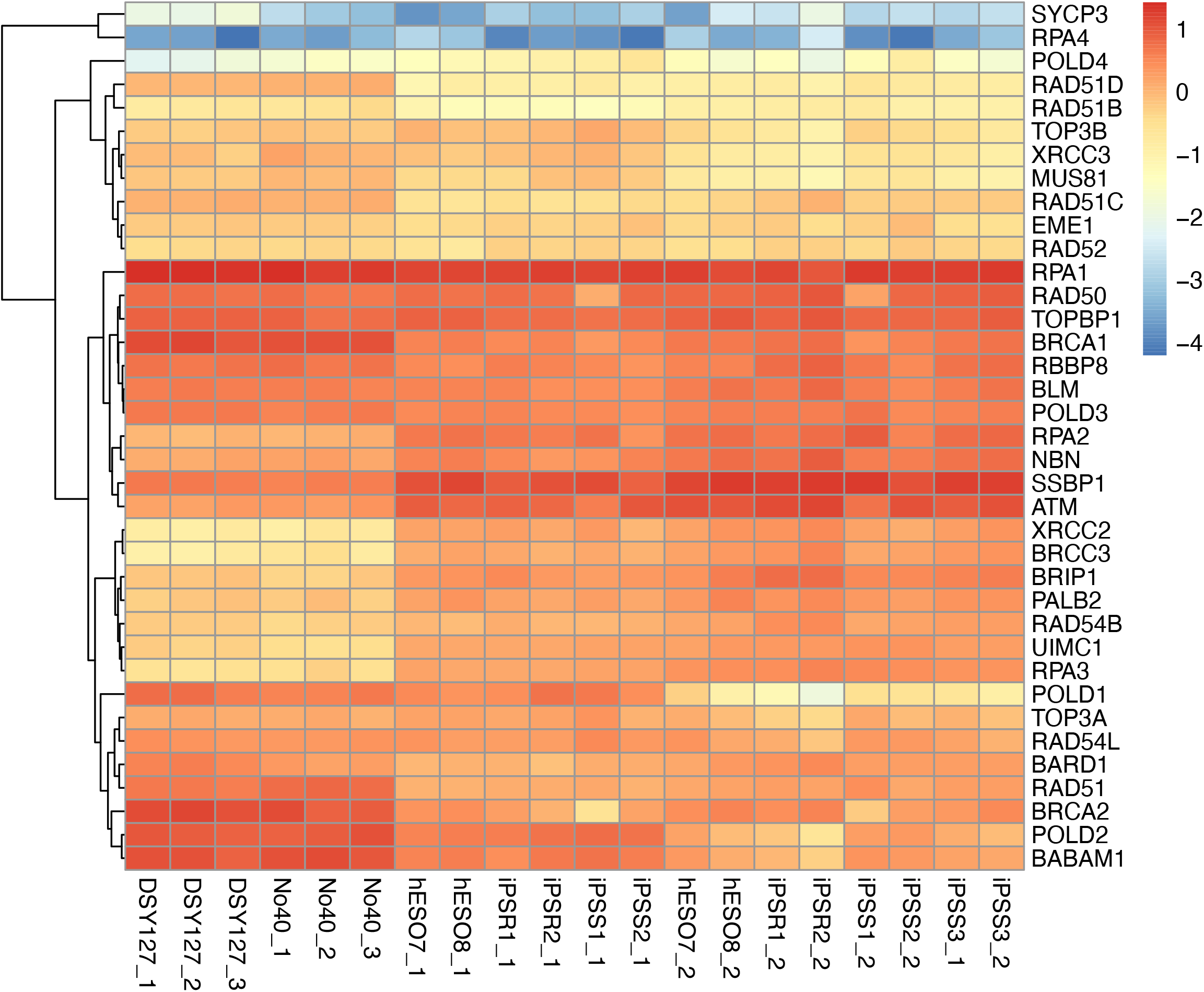
Normalized genes expression of HR-related genes in human and marmoset PSCs. The expression levels were treated with log2 and z-score normalization.

**Figure S3.**
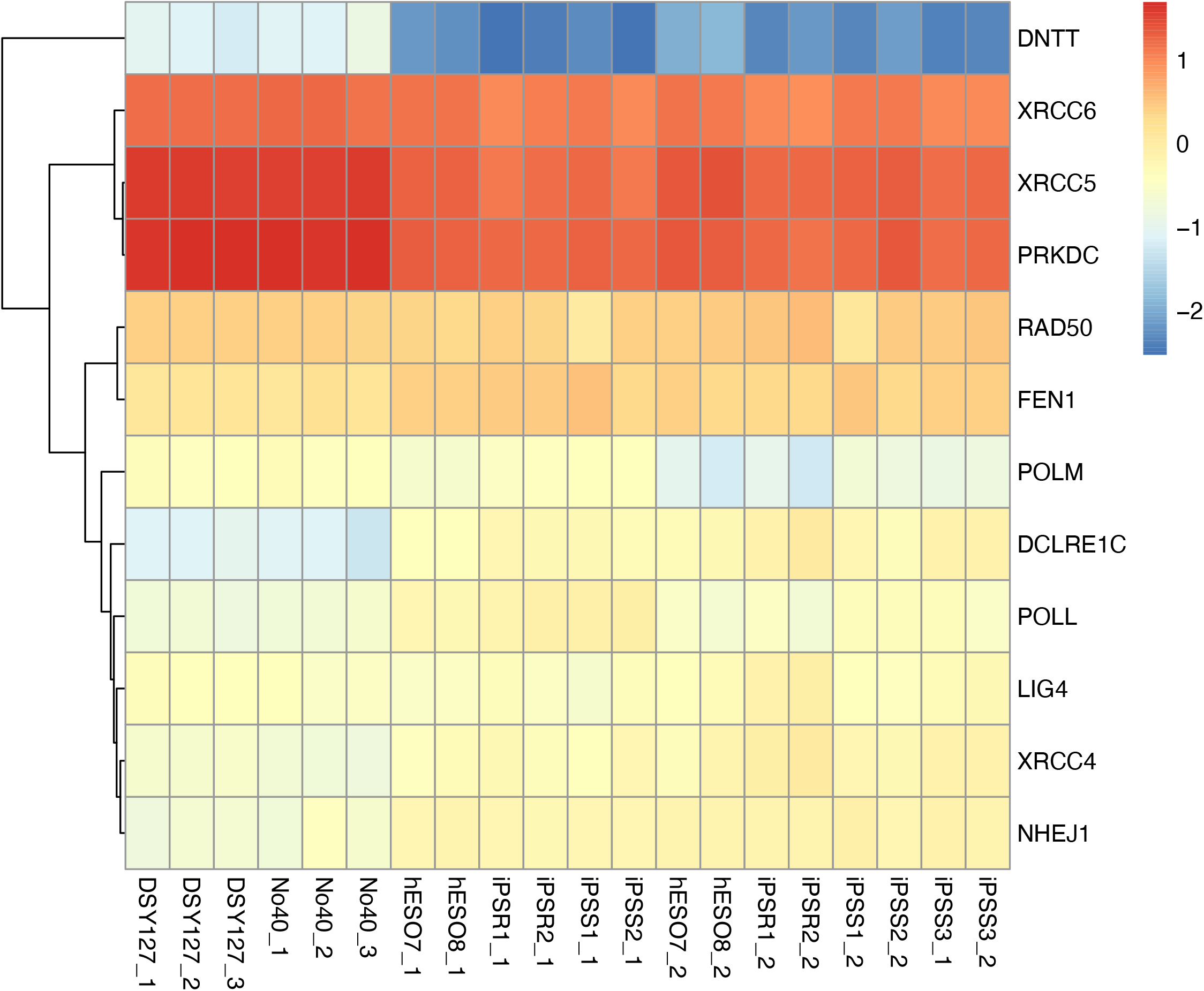
Normalized genes expression of NHEJ-related genes in human and marmoset PSCs. The expression levels were treated with log2 and z-score normalization.

**Figure S4.**
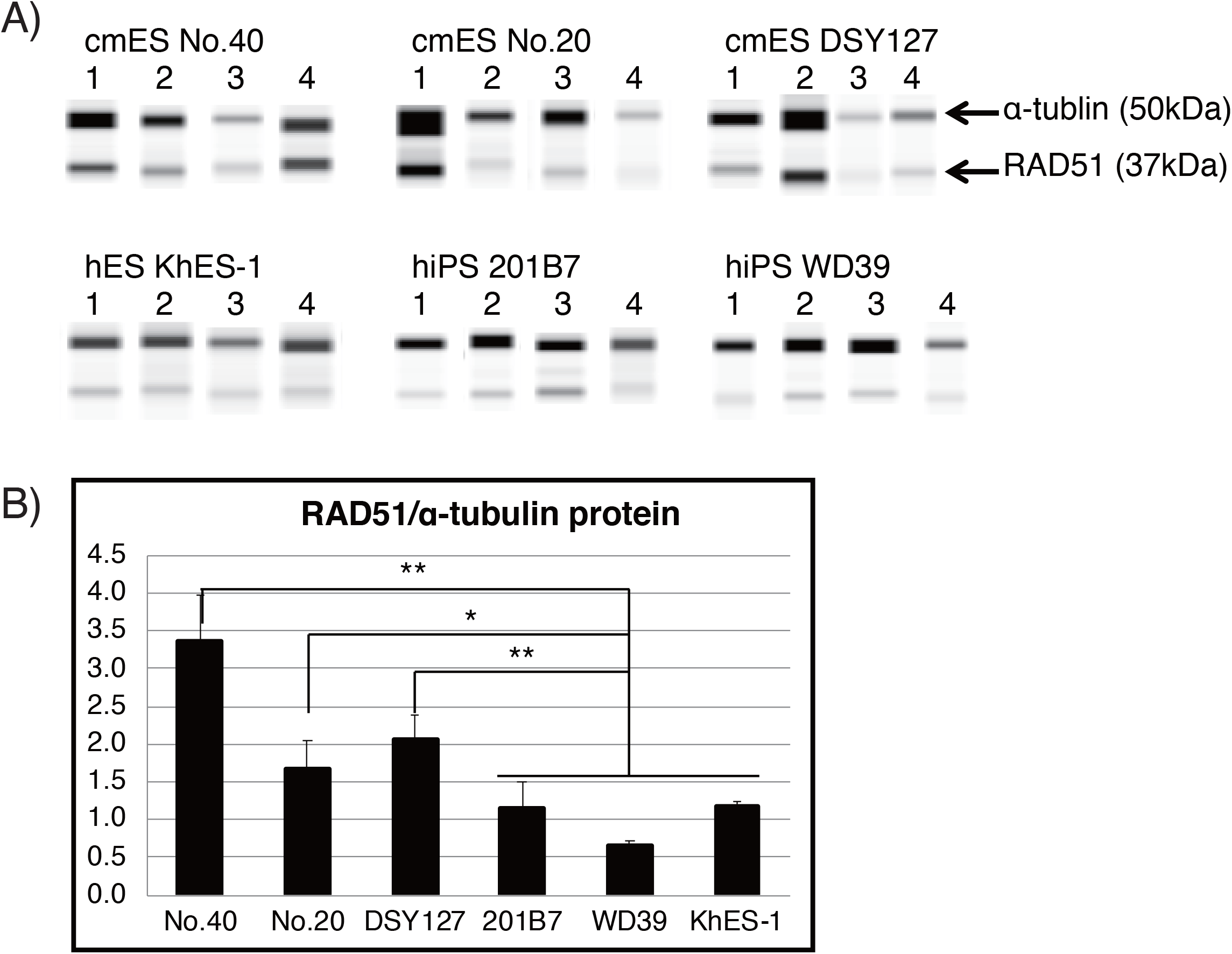
Western blotting analysis of RAD51 protein in human and marmoset PSCs. (**A**) Raw data of digital electrophoresis simultaneously detecting RAD51 (37 kDa) and α-tubulin (50 kDa). (**B**) Average RAD51 expression (normalized against α-tubulin) in each PSC line.

