## Supplementary Material and Supplementary Data 1 for "Comparative analyses of gene expression in common marmoset and human pluripotent stem cells (PSCs) identify factors enhancing homologous recombination efficiency in the *HPRT* locus of human PSCs": Supplementary Material.docx

**Supplementary Discussion**

*RAD51* and *DMC1* are the eukaryotic homologues of bacterial *RecA* and archaeal *RadA* (Chen et al., 2008; Lin et al., 2006) and are well-known key factors for the HR pathway (Baumann and West, 1998). The RAD51 protein forms a nucleoprotein filament on single-stranded DNA (ssDNA) to promote DNA-homology search, recognition, and strand exchange (Bianco et al., 1998) by exploiting two motifs critical for homology search and subsequent recombination (Ito et al., 2020). The overexpression of *RAD51* reportedly contributes to the enhancement of the HR ratio in multiple cell lines (reviewed in (Klein, 2008)), but there is considerable variation in the extent of this effect depending on experimental settings and cell lines. (Takayama et al., 2017), reported that *RAD51* overexpression in hPSCs did not result in a significant enhancement of the HR ratio in CRISPR-Cas9 mediated gene editing, but did cause an increase in biallelic KI. The latter might have resulted from the putative enhancement of RAD51 overexpression-induced inter-homolog repair (Wilde et al., 2018), classically known as sister-chromatid exchange (Painter, 1980; Sonoda et al., 1999). Our results on *RAD51* overexpression here were not fully consistent with the data from Takayama et al. (2017). This disparity might have resulted from the presence of different rate-limiting factors for HDR/NHEJ choice in different loci, cell types and lines.

From prokaryotes to eukaryotes, additional *RecA*-like genes have evolved following multiple duplication events and divergent evolution of the ancestral gene (Lin et al., 2006; Thacker, 1999). In vertebrate species, five genes have been identified as *RecA*/*RAD51* paralogues, namely, *RAD51B*, *RAD51C*, and *RAD51D* by homology searches (Albala et al., 1997; Dosanjh et al., 1998; Pittman et al., 1998), and *XRCC2* and *XRCC3* by their ability to complement susceptibility to ionizing radiation in the Chinese hamster CHO cell line (Cartwright et al., 1998; Tebbs et al., 1995). Structural studies revealed that the *RecA/RAD51* paralogues function mainly through formation of two distinct complexes (i) the RAD51C-XRCC3 complex, and (ii) the RAD51B-RAD51C-RAD51D-XRCC2 complex (Braybrooke et al., 2000; Liu et al., 2002). Respective knock-outs of these *RecA*/*RAD51* homologues in cell lines result in defective cell growth and HR, and hypersensitivity to DNA-damaging agents (Takata et al., 2001). Moreover, mice with mutations of the *RecA*/*RAD51* homologues show embryonic lethality (Kuznetsov et al., 2009; Thompson and Schild, 2001), indicating their importance for DNA repair, genomic stability, and cellular viability. On the other hand, the effect of overexpressing these paralogues has not been thoroughly investigated and nor has their expression patterns *in vivo*. In this context, our results showed, for the first time, that overexpressing these paralogues could enhance the efficiency of targeted gene engineering.

Heterozygous *BRCA1/2* mutations confer a high risk for breast and ovarian cancer in women (Ford et al., 1994; Futreal et al., 1994; Lancaster et al., 1996; Miki et al., 1994; Peto et al., 1999; Wooster et al., 1995). Moreover, cell lines and mouse mutants for these genes show various deficiencies, including hypersensitivity to irradiation, increased tumorigenesis, and HR deficiency; mouse embryos homozygous for *Brca1* and *Brca2* null mutations (*Brca1*^-/-^ and *Brca2*^-/-^) show developmental arrest during fetal development (Sharan et al., 1997; Shen et al., 1998). As tumorigenesis in patients carrying inherited heterozygous *BRCA1/2* mutations is triggered by deactivation of the wild-type allele, *BRCA1/2* are generally considered as tumor-suppressor genes.

The BRCA1 protein binds to many cellular proteins *in vivo*. For example, the BRCT domain in the C-terminus of BRCA1 recognizes a variety of DNA repair proteins (Deng and Brodie, 2000). The BRCA1 protein colocalizes with RAD51 in nuclear foci in mitotic cells (Scully et al., 1997); this colocalization is essential for HR upon replication fork stalling and collapse (Feng and Zhang, 2012). Additionally, via interaction with CtBP-interacting protein (CtIP), BRCA1 promotes ssDNA resection that is critical for NHEJ/HDR choice by competing with the HDR/MMEJ inhibitor protein P53BP1 (Chapman et al., 2012). Recent studies showed that knock-in (HDR) efficiencies with CRISPR-Cas9 were enhanced by direct or indirect suppression of P53BP1 activity (Jayavaradhan et al., 2019; Nambiar et al., 2019). Thus, the P53BP1-related mechanism provides a possible explanation for the ability of BRCA1 overexpression to enhance the HR ratio in hiPSCs.

The BRCA2 protein interacts directly with RAD51 through the BRC domain (Yang et al., 2005) and the C-terminus (Esashi et al., 2005). BRCA2 is required for recruitment of RAD51 to DSB sites, and is also important for its tumor-suppressive function (Moynahan et al., 2001). BRCA2 also has a DNA-binding domain that binds both ssDNA and dsDNA. These domains are thought to facilitate the formation of RAD51 nucleofilaments in HR. As a secondary role, BRCA2 may protect nascent DNA by MRE11A-mediated degradation upon replication fork stalling (Schlacher et al., 2011). These various properties clearly stress the importance of BRCA2 for HR and genomic stability; however, few studies have examined the effects of overexpression of *BRCA2* on HR. (Balia et al., 2011) reported that *BRCA2* overexpression, especially its cancer-associated variants, enhances spontaneous HR in HeLa cells; whether this effect occurs in other cell lines has not been thoroughly investigated. In this context, this study provides the first evidence that *BRCA2* overexpression in hPSCs has an HR-stimulating effect; this property might contribute to the development of stem cell-based technologies and therapies.
